# Systemic Brain Tumor Delivery of Synthetic Protein Nanoparticles for Glioblastoma Therapy

**DOI:** 10.1101/862581

**Authors:** Jason V. Gregory, Padma Kadiyala, Robert Doherty, Melissa Cadena, Samer Habeel, Erkki Ruoslahti, Pedro R. Lowenstein, Maria G. Castro, Joerg Lahann

## Abstract

Glioblastoma multiforme (GBM), the most aggressive form of brain cancer, has witnessed very little clinical progress over the last decades, in parts, due to the absence of effective drug delivery strategies. Intravenous injection is the least invasive delivery route to the brain, but has been severely limited by the blood-brain barrier (BBB). Inspired by the capacity of natural proteins and viral particulates to cross the BBB, we engineered a synthetic protein nanoparticle (SPNP) based on polymerized human serum albumin (HSA) equipped with the cell-penetrating peptide iRGD. SPNPs containing siRNA against Signal Transducer and Activation of Transcription 3 factor (STAT3*i*) result in *in vitro* and *in vivo* downregulation of STAT3, a central hub associated with GBM progression. When combined with the standard of care, ionized radiation, STAT3*i* SPNPs result in tumor regression and long-term survival in 87.5% of GBM bearing mice and primes the immune system to develop anti-GBM immunological memory.

---

Glioblastoma multiforme (GBM) is the most prevalent and aggressive form of brain cancer, currently accounting for approximately 47% of diagnosed brain cancers.^1^ GBM is characterized by its high invasiveness, poor clinical prognosis, high mortality rates and frequent recurrence.^2^ Current therapeutic approaches include focal radiotherapy, and adjuvant chemotherapeutics in combination with surgical resection. However, due to the delicate anatomical structure of the brain and the highly invasive nature of glioma cells, complete surgical resection is rarely achieved.^3^ Residual tumor cells infiltrate the surrounding brain tissue and are protected by the blood-brain barrier (BBB), rendering them unresponsive to conventional chemotherapeutic drugs.^4^ Despite recent surgical and chemotherapeutic advances, the median survival (MS) for patients diagnosed with GBM remains just 12-15 months with a 5-year survival rate of 5%. The development of alternative and novel delivery approaches to effectively treat GBM remains one of the most pressing challenges in cancer therapy.^5^

A growing body of evidence suggests that the Signal Transducer and Activator of Transcription 3 (STAT3) pathway is involved in multiple signaling pathways related to tumor progression and evasion of the immune system.^6–8^ Multiple growth factors and cytokines frequently overexpressed in cancer, such as EGF, FGF and IL-6, activate STAT3 via tyrosine phosphorylation.^8–10^ Activated STAT3 (pSTAT3) translocates to the nucleus and participates in the transcription of genes that inhibit apoptosis, and promote tumor cell proliferation and metastasis. Histopathological analysis of brain tumors demonstrated STAT3 to be overexpressed in patients with grade III astrocytomas and grade IV GBMs; increased STAT3 levels are negatively associated with MS in these patients.^7^ In previous studies, the STAT3 inhibitor CPA-7 induced tumor cell death in GL26 mouse GBM and HF2303 human GBM stem cells. However, a positive therapeutic effect was only observed in peripheral tumors, but not in intracranial tumors, pointing towards the inability of CPA-7, like many small molecule therapeutics and biomolecules, to cross the protective BBB and enter the brain tumor compartment.^6^

A wide range of nanoparticles have been developed to deliver chemotherapeutic drugs, such as docetaxel,^11–13^ paclitaxel,^14–16^ doxorubicin^17^ or other small molecule chemotherapeutics;^18–22^ or, to leverage antibodies,^23,24^ RNA,^25–28^ or peptides^29^ in an attempt to enhance GBM therapy. Despite these grand efforts, research conducted over the past decades has made only marginal advances with no real promise of a clinical path towards a viable curative treatment. In general, these nanocarriers share a number of common characteristics, e.g., (i) they are made of synthetic materials, (ii) they tend to accumulate and persist in liver and spleen causing severe side effects, and (iii) they are incapable of passing the BBB. In contrast, natural evolution has resulted in proteins and viral particulates that can target to and transport through the BBB.^30^ Inspired by the unique capabilities of biological nanoparticles, we engineered a GBM-targeting synthetic protein nanoparticle (SPNP) comprised of polymerized human serum albumin (HSA) and oligo(ethylene glycol) (OEG), loaded with the cell-penetrating peptide iRGD^31–33^ as well as STAT3*i*. The choice of HSA as the major matrix component was motivated by its rapid and well-understood clearance mechanisms, its demonstrated clinical relevance, and its exquisite biochemical compatibility with both, therapeutic agents and homing peptides. In addition, albumin-based nanocarriers have been shown to engage cell-surface receptors, such as SPARC^34^ and gp60,^35^ that are overexpressed on glioma cells and tumor vessel endothelium.^36–38^

## Particle Design, Synthesis and Characterization

SPNPs were prepared via electrohydrodynamic (EHD) jetting, a process that utilizes atomization of dilute solutions of polymers to produce well-defined nanoparticles.^39–41^ Rapid acceleration of a viscoelastic jet in an electric field leads to a size reduction by several orders of magnitude facilitating rapid solvent evaporation and solidification of the non-volatile components into nanoparticles. Here, the jetting solution was comprised of human serum albumin and a bifunctional OEG macromer (NHS-OEG-NHS, 2kDa), which were mixed with various additional components, such as a therapeutic siRNA, polyethyleneimine (PEI, a siRNA complexing agent), and the tumor penetrating peptide, iRGD, prior to nanoparticle preparation. Similar to a step-growth polymerization, the OEG macromer combined with albumin molecules through reaction with its lysine residues resulting in water-stable nanoparticles. After EHD jetting and collection, the resulting SPNPs had an average diameter of 115 ± 23 nm in their dry state (Fig. 1b). Once fully hydrated, the average diameter of SPNPs increased to 220 ± 26 nm based on dynamic light scattering (DLS) measurements (Supplementary Fig. 1). The degree of nanoparticle swelling was controlled by varying the HSA-to-OEG ratios between 4:1 and 20:1 and the molecular weight of the OEG macromer between 1 kDa and 20 kDa. An increase of the OEG content from 5% to 20% resulted in a reduction of particle swelling by 20 volume-%. SPNPs were stable for at least 10 days at 37 °C under physiological conditions; with no significant change in particle size or morphology (Supplementary Fig. 2). When exposed to mildly acidic conditions (pH 5.0), similar to those observed in endosomes of cancer cells, the diameters of SPNPs increased to 295 ± 31 nm (Fig. 1c). We note that defining particle properties, such as particle size, shape and swelling behavior, was, within the margins of error, identical for fully loaded SPNPs, empty nanoparticles and nanoparticles loaded with siRNA and/or iRGD.

**Fig. 1:**
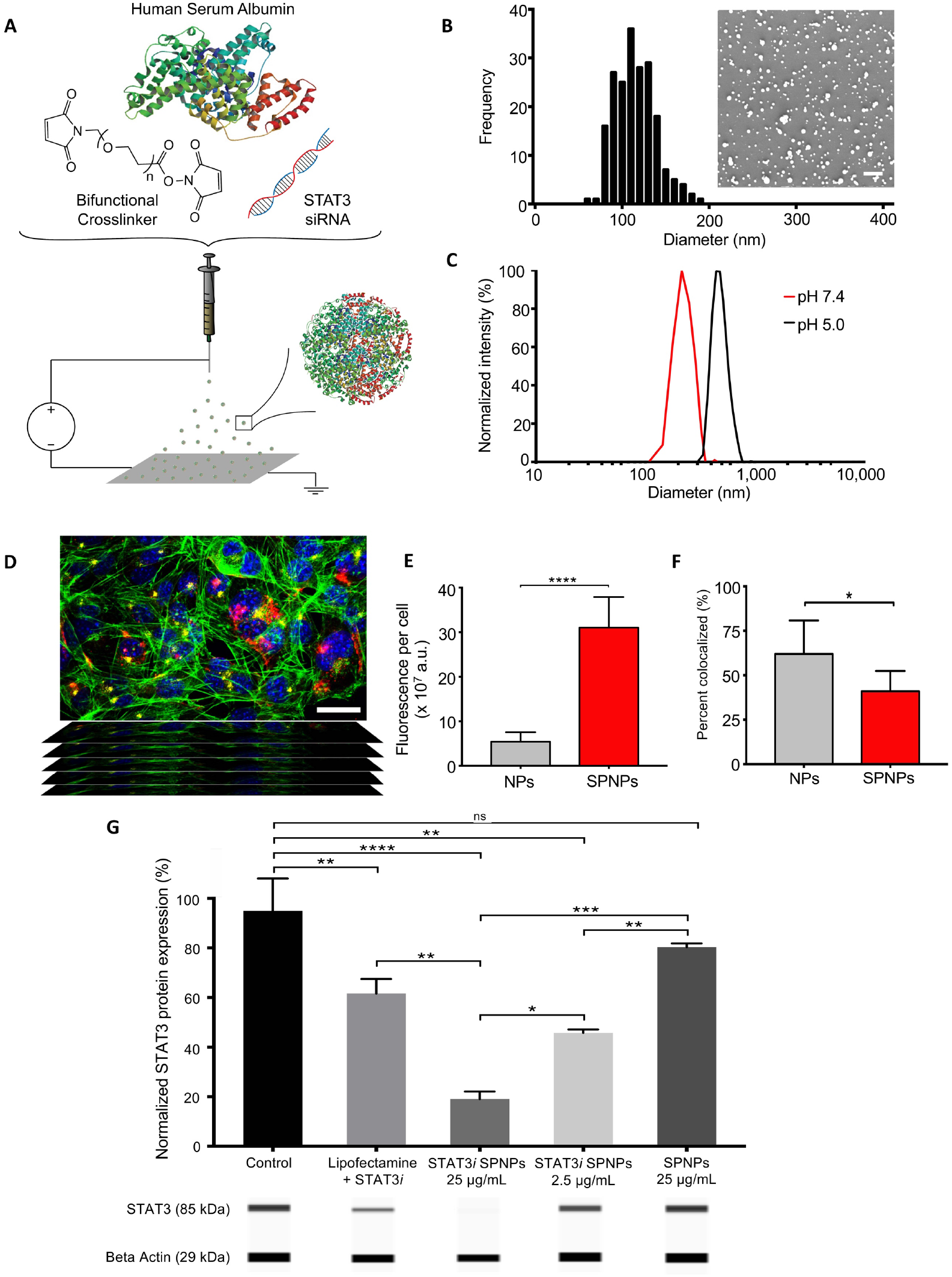
STAT3 expression is effectively silenced in vitro by siRNA-loaded SPNPs. **(A)** Schematic of jetting formulation for crosslinked, STAT3i-loaded, iRGD-conjugated, targeted albumin nanoparticles (STAT3i SPNPs) **(B)** Particle size characterization and analysis was performed using Scanning Electron Microscopy (SEM). Average particle diameter, 115 ± 23.4 nm. Scale bar 1 µm. **(C)** Particles undergo swelling in their hydrated state and further swell at reduced pH. Average diameters: pH 7.4, 220 ± 26.1 nm. pH 5.0, 295 ± 31.2 nm. **(D)** Representative confocal z-stack image of cells seeded in the presence of SPNPs **(E)** Local release of iRGD from SPNPs increases particle uptake in GL26 glioma cells by greater than five-fold. **(F)** Internalized SPNPs colocalize with lysosomes to a lesser extent than untargeted particles. **(G)** STAT3 siRNA-loaded SPNPs significantly reduce *in vitro* expression of target protein in GL26 glioma cells compared to untreated and empty particle control groups. Bars represent mean ± standard error, (****p<0.0001, ***p<0.0005, **p<0.005, *p<0.05, two-way ANOVA test) (n ≥ 3 wells/treatment group).

## In Vitro Cell Uptake and siRNA Activity

Previously, co-delivery of the cell-penetrating peptide iRGD has increased tumor targeting for both, small drugs and iron oxide nanoparticles.^42^ The iRGD peptide has been shown to act in a three-stage process, first binding to αvβ3 and αvβ5 integrins, followed by a proteolytic cleavage step and a secondary binding event to neurophilin-1 (NRP-1), which activates an endocytotic/exocytotic transport pathway.^43^ In the past, iRGD mediated tumor homing has been approached either in form of systemic co-delivery of iRGD with nanoparticles^44,45^ or by decorating the nanoparticles with surface-bound iRGD.^46,47^ In our case, the iRGD peptide is preloaded into SPNPs during EHD jetting to promote local release at the vicinity of the BBB. To investigate the intracellular fate of nanoparticles and the effect of iRGD as a targeting ligand, in vitro uptake experiments were performed. When iRGD-loaded SPNPs were incubated with GL26 glioma cells for a period of 30 minutes, particle uptake increased by ~ 5-fold compared to nanoparticles without iRGD (Fig. 1e). Image analysis of 3D confocal images (Fig. 1d) using CellProfiler revealed that significantly less particles colocalized with lysosomes of GL26 glioma cells compared to nanoparticles without iRGD (Fig. 1f). These data suggest that the delivery of iRGD from SPNPs results in higher uptake and higher cytosolic nanoparticle concentrations. The above-mentioned pH-dependent swelling of SPNPs, along with the “proton sponge” effect previously postulated for PEI,^48^ may contribute to more effective particle escape from endocytotic vesicles, enhancing overall siRNA delivery to the cytosol and RNA-induced silencing.^49,50^

We next evaluated if siRNA loaded into SPNPs during EHD jetting remains biologically active. First, siRNA loading and release from SPNPs was evaluated using a Cy3-labeled STAT3*i* surrogate. Utilizing stimulated emission depletion (STED) microscopy, we confirmed uniform distribution of siRNA throughout the entire NP volume (Supplementary Fig 3). *In vitro* release of fluorescently tagged siRNA confirmed that 96% of the initial amount of siRNA was encapsulated into SPNPs; corresponding to a siRNA loading of 340 ng, or 25 pmol of siRNA per mg of SPNPs. Furthermore, we observed that approximately 60% of the encapsulated siRNA was released over the first 96 hours, followed by a sustained release period progressing for 21 days (Supplementary Fig. 4). When albumin NPs were loaded with siRNA against GFP, SPNPs significantly suppressed GFP expression in mouse glioma cells transfected to express mCitrine (GL26-Cit, Supplementary Fig. 5) relative to control albumin NPs loaded with scrambled siRNA or free GFP siRNA that was delivered using lipofectamine as the transfection agent. Moreover, protein knockdown persisted significantly longer in the SPNP group than in lipofectamine-transfected cells (Supplementary Fig. 5). While the latter entered a recovery phase after two days and nearly returned to normal GFP levels by day five, cells treated with GPF*i* SPNPs showed sustained protein knockdown throughout the experiment. There were no significant differences in particle size, surface charge, or morphology between siRNA-loaded SPNPs and the control particles.

For particle concentrations of 2.5 and 25 ug/mL, SPNPs co-loaded with iRGD and STAT3*i* significantly reduced total STAT3 protein expression relative to the untreated control group or empty SPNPs (Fig. 1g). Moreover, we observed a dose-dependent response in that a higher particle concentration resulted in ~ 2-fold further decrease in total STAT3 expression. No detectable signs of cytotoxicity were observed for any of the tested NP groups, which we attributed to the fact that the delivered siRNA concentrations were below the cytotoxicity limit observed for free STAT3 siRNA in GL26 cells (Supplementary Fig. 6). Based on these *in vitro* experiments, we chose an effective dose of 5 μg/kg in subsequent animal studies.

## Systemic Delivery of SPNPs in an Intracranial GBM Model

In the past, the BBB has been an unsurmountable delivery challenge for nanocarriers^51,52^ that are systemically administrated via IV injection. To evaluate, if systemically delivered SPNPs can home to brain tumors, SPNPs loaded with Alexa Fluor™ 647-labeled albumin were prepared as described above. In the absence of large animal GBM models, we selected the very aggressive GL26 syngeneic mouse glioma model, which is known to exhibit histopathological characteristics encountered in human GBM,^53^ to evaluate GBM-targeting of SPNPs. In addition, this model features an uncompromised immune system, which was deemed to be essential, because of the prominent role that STAT3 plays in downregulation of the immune system. A dose of 2.0 × 10^13^ SPNPs was delivered to GBM-bearing mice via a single tail vein injection seven days after glioma cell (GL26-Citrine) implantation in the right striatum of the mice (Fig. 2a). After 4 or 24 hrs, animals were euthanized, and brains were collected, sectioned, and stained prior to confocal imaging. While the majority of particles was taken up by other organs, such as liver, kidney, spleen and the lungs, a significant number of SPNPs appeared to have crossed the BBB and were identified within the brain tumor microenvironment at both time points (Fig. 2b). Tumor targeting was markedly increased after 24 hrs, hinting towards the possibility that secondary transport processes, such as transcytotic or immune-cell-mediated BBB pathways, are contributing to the brain-homing of SPNPs, in addition to the direct targeting of the brain endothelium by circulating nanoparticles. The notion of a multivariant transport mechanism is consistent with our finding that SPNPs were localized inside of tumor cells (green) and macrophages (red), suggesting that both cell types can internalize SPNPs (Fig. 2b).

**Fig. 2:**
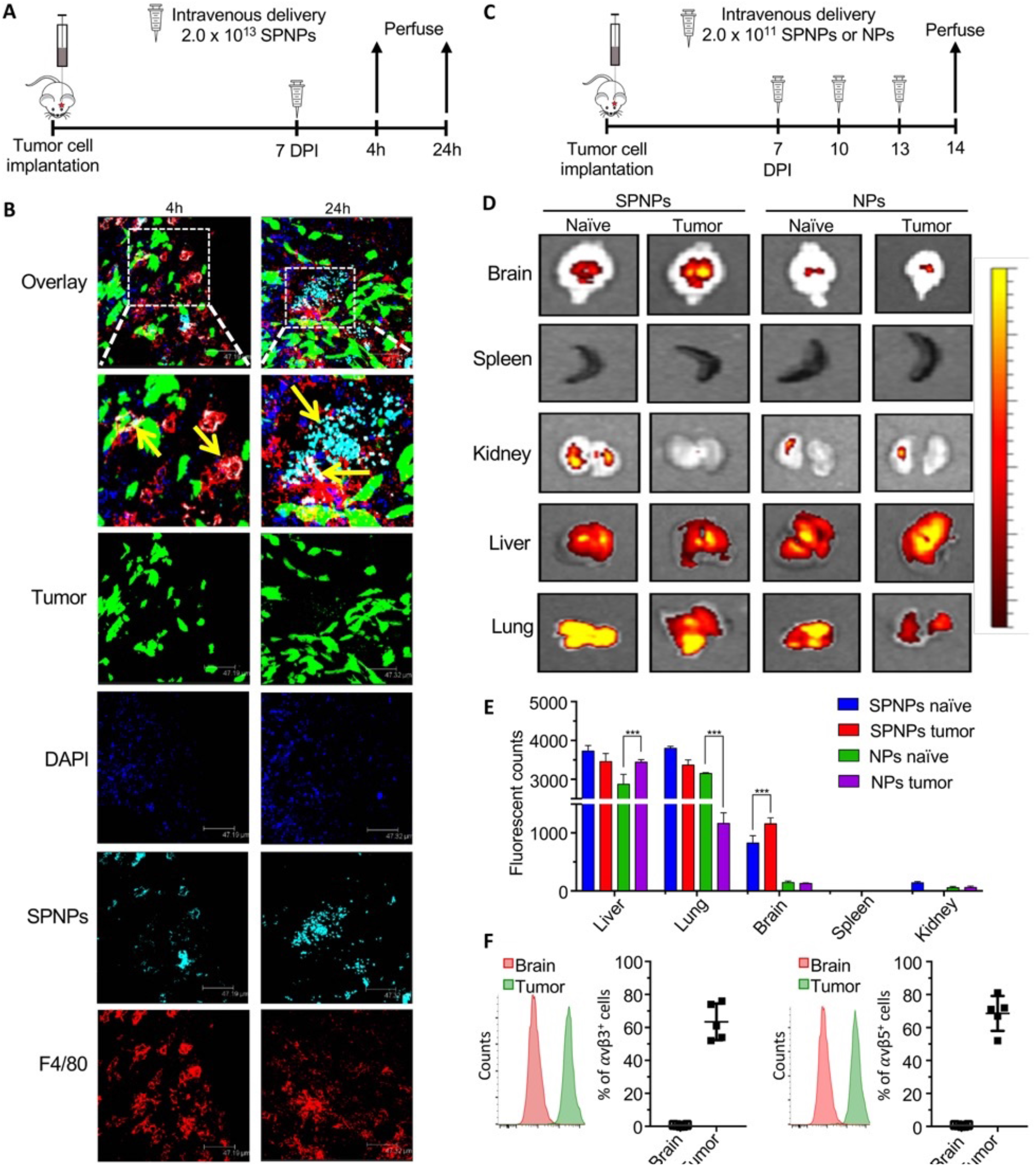
In vivo brain targeting and biodistribution of SPNPs. **(A)** Timeline for the tumor targeting study. Mice were IV administrated a single 2.0 × 10^13^ SPNPs or Empty-NPs dose via the tail vein seven days post GL26 tumor cells implantation. Confocal imaging of sectioned brains was performed 4 and 24 hours post particle administration. **(B)** Alexa Fluor 647 labeled SPNPs (cyan) colocalize (indicated with yellow arrows) with macrophages (red) and tumor cells (green, mCitrine). Less NPs are observed in the tumor microenvironment 4h post systemic delivery compared to 24h. **(C)** Timeline representation of the biodistribution study. Mice were IV administered 2.0 × 10^11^ SPNPs or Empty-NPs 7, 10, and 13 days post tumor cell implantation or saline injection. **(D)** Fluorescence imaging of tumor-naive and tumor-bearing mice organs sacrificed at 24h post final NP delivery. **(E)** Quantitative analysis of NP biodistribution within the tumor and peripheral organs (****p < 0.00001, **p < 0.001, *p < 0.05, two-way ANOVA test) **(F)** Quantitative flow cytometry results of avβ3 and avβ5 integrin expression in normal brain tissue and GL26 tumors.

Using the same intracranial tumor model, GBM specific biodistribution of SPNPs was assessed. Tumor-bearing mice were injected three times (7d, 10d, and 13d DPI) with Alexa Fluor 647-labeled SPNPs or nanoparticles without iRGD (Fig. 2c). In addition, naïve mice (non-tumor bearing) were subjected to the same regimen to delineate tumor-specific characteristics. After 14 days, naïve and tumor bearing mice were euthanized, their main organs were collected, and the nanoparticle distribution was analyzed via fluorescence imaging (Fig. 2d). In both GBM-bearing and naïve mice, significantly more SPNPs were observed in the brain compared to nanoparticles without iRGD. As expected, SPNPs also accumulated in the lungs and liver – independent of the particular experimental group. The former can be attributed to being the first capillary bed the NPs would encounter following intravenous injection, while the latter represents the primary route of clearance for NPs of this size.^54^ Brain accumulation of SPNPs loaded with iRGD was higher for both naïve and GBM-bearing mice compared to iRGD-free nanoparticle groups. When directly comparing SPNP localization within the brain compartment, the accumulation of iRGD-loaded SPNPs was 40% higher in tumor-bearing brains (Fig. 2e) than in healthy brains.

To provide a rationale for the increased accumulation of SPNPs in the brains of tumor-bearing mice, we searched for integrin-expressing cells in the tumor microenvironment. Specifically, we focused on the relative expression of αvβ3 and αvβ5 integrins, because these ligands have been shown to play pivotal roles in the iRGD-induced accumulation of nanoparticles and small drugs in tumors,^31^ and are overexpressed in gliomas.^32^ Using the GBM model and dosing schedule from the biodistribution studies, brains from GBM tumor-bearing mice were collected at 23 DPI. Normal brain and tumor tissue were dissected from the brain, processed, and stained with αvβ3 and αvβ5 antibodies for flow cytometry analysis. More than 60% of the GBM tumor population expressed αvβ3 and αvβ5 integrins at high levels, while normal brain cells showed minimal expression of these proteins. These results, along with the observed differences in brain accumulation in the biodistribution study, appear to be consistent with the previously postulated hypothesis that iRGD promotes local brain tumor accumulation.^43^

## *In vivo* Survival Studies

To test the efficacy of SPNPs *in vivo*, GBM-bearing mice were treated intravenously with multiple doses of STAT3*i* SPNPs over the course of a three-week treatment regimen (Supplemental Fig. 9). After tumor implantation, the median survival (MS) of untreated mice was about 28 days. In mice that received multiple doses of empty SPNPs, the MS remained unaltered (28 days). In contrast, when SPNPs loaded with STAT3*i* were administered, the MS increased to 41 days, a statistically significant increase of 45%. Delivery of the same doses of free STAT3*i* resulted in a modest extension of MS by 5 days, which is likely too low to elicit a significant therapeutic effect. The low efficacy of free STAT3*i* can be explained by the rapid degradation of genetic material following systemic administration – in addition to siRNA’s inability to cross the BBB.^55^

Encouraged by the prospect of a nanoparticle formulation for STAT3*i* delivery with significant *in vivo* efficacy, we combined STAT3*i* SPNPs with the current standard of care, i.e., focused radiotherapy (IR). Previous studies have identified a direct correlation between STAT3 overexpression and radioresistance in other cancers,^56,57^ suggesting that its knockdown could contribute to enhanced efficacy. We thus established a treatment protocol that combined the previously evaluated multi-dose regimen with a repetitive, two-week focused radiotherapy regimen (Fig. 3a). Once GBM tumors had formed, mice received seven doses of STAT3*i* SPNPs over the course of a three-week period. During each of the first two weeks, mice also received five daily 2 Gy doses of IR for a total of ten treatments (Fig. 3a). Experimental groups included mice that received either STAT3*i* SPNPs, empty SPNPs, free STAT3*i*, or saline with or without combined radiotherapy. In all cases, the addition of radiotherapy increased the MS, with IR alone resulting in a MS extension from 28 to 44 DPI (Fig. 3b). Combining IR with empty SPNPs did not further alter the MS. Consistent with our previous experiment, free siRNA provided a slight, statistically significant benefit, where the MS was increased to 58 DPI, when combined with IR (Fig. 3b, brown line). However, a more significant effect was observed for the combination of STAT3*i* SPNPs with IR. Of the eight mice in this group, seven reached the standard long-term survivor time point of 90 DPI and appeared to be completely tumor-free thereafter (Fig. 3b, blue line). The single mouse receiving this treatment that did not reach long-term survival was moribund at 67 DPI, living longer than any other non-surviving subject from all other groups.

**Fig. 3:**
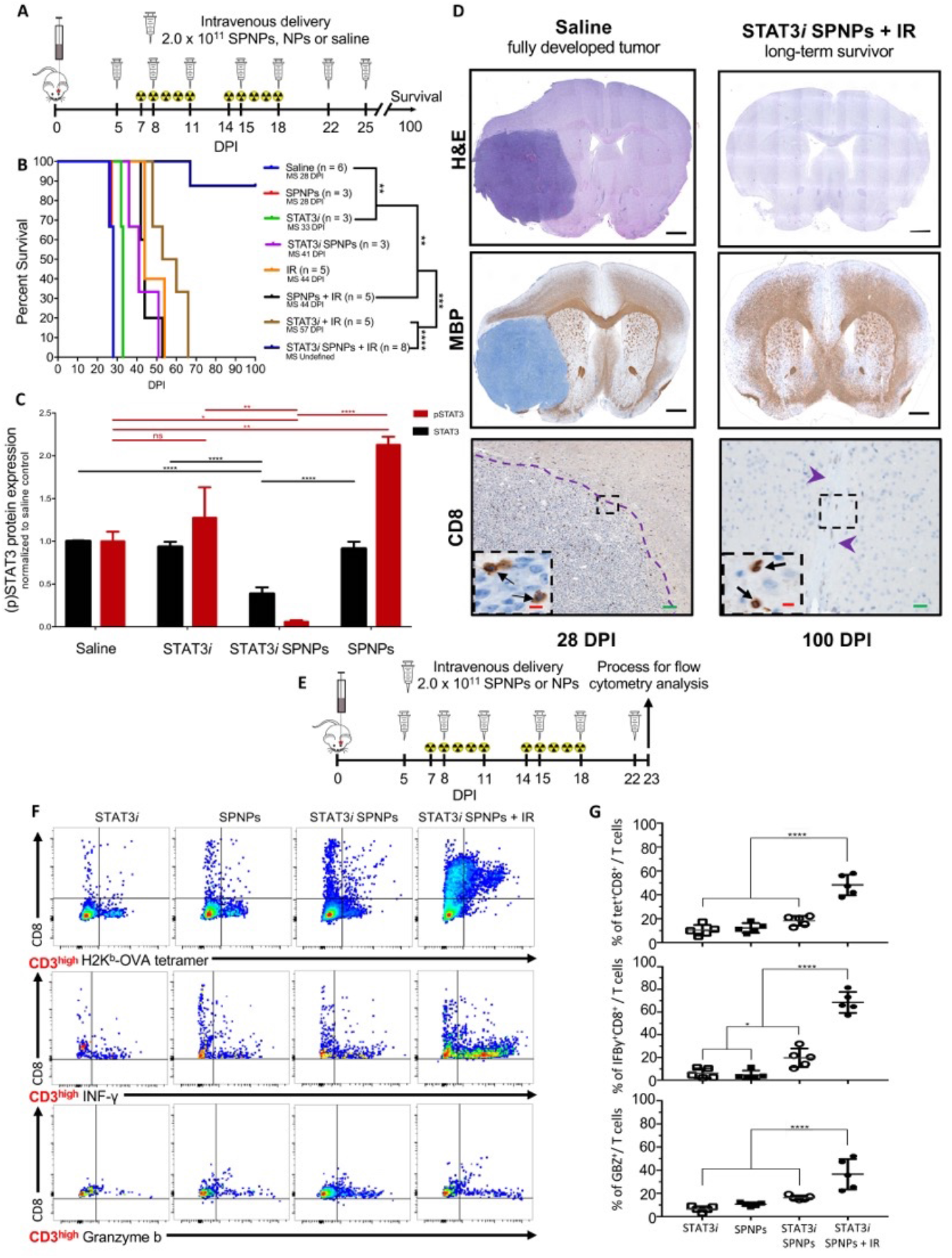
Systemic delivery of STAT3i SPNPs in combination with radiotherapy (IR) results in increased survival and primes an adaptive immune response. **(A)** Timeline of treatment for the combined NP + IR survival study. **(B)** Kaplan-Meier survival curve. Significant increase in median survival is observed in all groups receiving IR. Mice (7/8) treated with STAT3*i* SPNPs + IR reach long-term survival timepoint (100 DPI) with no signs of residual tumor. **(C)** Quantified STAT3 expression for resected brains from the survival study, brains were collected when mice displayed signs of neurological deficits. A significant reduction is STAT3 and pSTAT3 expression was observed in the STAT3*i* SPNP cohort relative to untreated control. Both soluble STAT3*i* and empty SPNPs (with no siRNA) did not have high total STAT3 expression but they had increased levels of active phosphorylated STAT3 expression (pSTAT3). Bars represent the mean relative to untreated control ± standard error (n ≥ 3). ****p < 0.0001, **p < 0.005, *p < 0.05,; unpaired t-test. **(D)** IHC staining for untreated control and STAT3*i* SPNP + IR long-term survivor. (Top) H&E staining shows the fully formed tumor in the saline control group (28 DPI). When treated with the combination of STAT3*i* SPNPs + IR, no tumor or signs of necrosis were observed. (Middle) MBP staining shows preserved brain structures with no apparent changes in oligodendrocyte integrity in mice that received STAT3*i* SPNPs + IR treatment compared to the saline control. (Bottom) CD8 staining shows no overt inflammation in mice that received STAT3*i* SPNPs + IR treatment compared to the saline control. **(E)** Timeline of TME immune population by flow cytometry. **(F)** Flow cytometry analysis of CD8 cells in the TME. Representative flow plots for each group are displayed. **(G)** Quantitative analysis of tumor-specific CD8^+^ T cells within the TME. GL26-OVA tumors were analyzed by staining for the SIINFEKL-K^b^ tetramer. Activation status of CD8^+^ T cells within the TME was analyzed by staining for granzyme B (Gzb) and IFNγ after stimulation with the tumor lysate.. ****p < 0.0001; unpaired t-test. Bars represent mean ± SEM (n = 5 biological replicates).

In order to characterize the effects of the combined treatments, additional studies were performed. Following the same treatment outlined in Fig. 3a, we elucidated the expression of STAT3 and its active phosphorylated form, pSTAT3, in the brain tissues (Fig. 3c). As expected, the greatest reduction in both the total and phosphorylated form of the protein were found in the STAT3*i* SPNP group. Greater than 50% reduction in total STAT3 protein was present, compared to the saline treated control. Phosphorylated STAT3 levels were even more dramatically reduced, with a greater than ten-fold reduction in pSTAT3 levels comparing the same groups. In contrast, the total STAT3 levels were relatively unchanged in both, the free STAT3*i* and empty SPNP groups, compared to untreated control. Here, pSTAT3 was increased by 110% in the cohort receiving empty SPNPs, suggesting a shift in the balance of the two protein forms, perhaps due to a localized upregulation of kinase activity.

Immunohistochemistry (IHC) analysis was used to compare the brains from long-term survivors to other treatment groups. Fig. 3d shows a direct comparison of the brain of a long-term survivor and that of a control animal that did not receive treatment. Hematoxylin and eosin (H&E) staining clearly shows the presence of a fully developed tumor localized within a single hemisphere of the saline treated mice. Conversely, we observe no overt signs of tumor presence in the STAT3*i* SPNP + IR treated survivors (Fig. 3d, top). Moreover, no apparent regions of necrosis, palisades or hemorrhages were present in these animals 90 DPI after receiving a full course of therapy. Myelin basic protein (MBP) staining was performed to assess the integrity of oligodendrocytes, an indicator for the disruption of surrounding brain architecture. No apparent changes in oligodendrocyte integrity was observed in mice that received the combined STAT3*i* SPNP + IR treatment when compared to the cancer-free right faces of mice in the saline treated control group (Fig. 3d, middle). In addition, CD8 T cells were sparse in the TME (Fig. 3d, bottom) and their total number was significantly reduced in STAT3*i* SPNP + IR treated mice compared to the saline treated control group (Supplementary Fig. 10). Independent pathological analysis of potential side effects affecting the livers of mice treated with STAT3*i* SPNP + IR therapy found minimal to mild monoculear pericholangitis across all groups and characterized it as spontaneous background rather than a direct result of the applied therapy. (Supplementary Fig. 11). In all treatment groups, with the exception of the saline treated control, minimal to mild coagulative necrosis was present. In the treatment group that received free STAT3*i*, one animal displayed multiple foci of coagulative necrosis, which distinguished it from all other animals in the entire study cohort, including those from the combined STAT3*i* SPNP + IR group, where the regions of necrosis were generally small and were deemed not to induce a biologically significant effect on liver function.

Next, SPNP-induced immune responses were assessed using a modified enzyme-linked immunosorbent assay (ELISA, Supplementary Fig. 12). To avoid a species-to-species mismatch due to the use of human serum albumin in mice, otherwise identical NPs were synthesized, in which HSA was replaced with mouse serum albumin (MSA). No circulating antibodies specific to MSA SPNPs were observed in any of the treatment groups (Supplementary Fig. 12) indicating neglectable immunogenicity against any of the individual components of SPNPs, such as OEG, STAT3*i*, iRGD, or PEI. As expected, replacing MSA with HSA resulted in elevated levels of HSA antibodies for both, STAT3*i* SPNP and empty SPNP treatment groups (Supplementary Fig. 12). However, free STAT3*i* therapy did not induce this same response suggesting that antibodies were formed in response to the exposure of the nanoparticles rather than the active therapeutic ingredient.

To further investigate the potential role of the adaptive immune system, we more closely examined the population of CD8 T cells within the tumor microenvironment (TME) via flow cytometry. We established tumors in mice using GBM cells that harbored a surrogate tumor antigen ovalbumin and compared the responses elicited by the various treatment formulations (Fig 3e). Tumor specific T cells in the TME were characterized by staining for the SIINFEKL-H2Kb-OVA tetramer, an OVA cognate antigen within the CD8 T cell population. Tumor specific CD8 T cells (CD3 + / CD8 + /SIINFEKL-H2K b tetramer) within the STAT3*i* SPNP + IR group were increased by twofold compared to all other groups (Fig. 3f, top). Staining the same population of cells with INF-γ and granzyme B (GZB) revealed a twofold increase in cytotoxic T cells in the TME (Fig. 3f, middle and bottom) in the STAT3*i* SPNP + IR group relative to all other groups. Taken together, these results suggest a robust anti-GBM response elicited by the combined NP + IR therapy that is likely contributing to the observed therapeutic success.

Flow cytometry analysis of tumor infiltrating macrophages and conventional dendritic cells (cDCs:CD45 + /CD11c + / B220 −) was used to compare treatment groups containing free STAT3*i*, empty SPNP and STAT3*i* SPNPs in combination with IR (Fig. 4a and 4b). Co-staining of CD45^+^ cells with F4/80 and CD206 antibodies was used to establish a subpopulation of tumor associated macrophages (TAMs). Within the TAM population, both, M1 (CD45 + /F4/80 + / CD206 −) and M2 (CD45 + /F4/80 + / CD206 +) macrophage phenotypes, were identified for all cohorts, but their relative abundance was significantly different in the STAT3*i* SPNP + IR group compared to all other groups. In the STAT3*i* SPNP + IR group, M1 macrophages were increased by 2.5-fold (Fig. 4c, left), whereas the number of M2 macrophages was decreased by three- to four-fold decrease (Fig. 4c, middle). These findings are consistent with the notion that the STAT3*i* SPNP + IR treatment selectively decreases the immune suppressive M2 macrophage subpopulation. In addition, antigen presentation by cDCs was significantly higher in animals receiving STAT3*i* SPNPs compared to free siRNA and empty SPNPs (Fig. 4c, right). IR treatment further elevated this effect resulting in the largest cDC population for the STAT3*i* SPNP + IR group. Co-staining of CD45 and F4/80^high^ cells with CD206 and Arg1 antibodies in the STAT3*i* SPNP + IR group confirmed that the vast majority of TAMs were of the M2 phenotype (Supplemental Fig. 13). Among all TME CD45^+^ immune cells, only M2 macrophages displayed the far-red Alexa Fluor 647 signal indicative of SPNPs suggesting that immune suppressive M2 macrophages are the primary TME-based immune cells that internalize SPNPs (Supplemental Fig. 14).

**Fig. 4:**
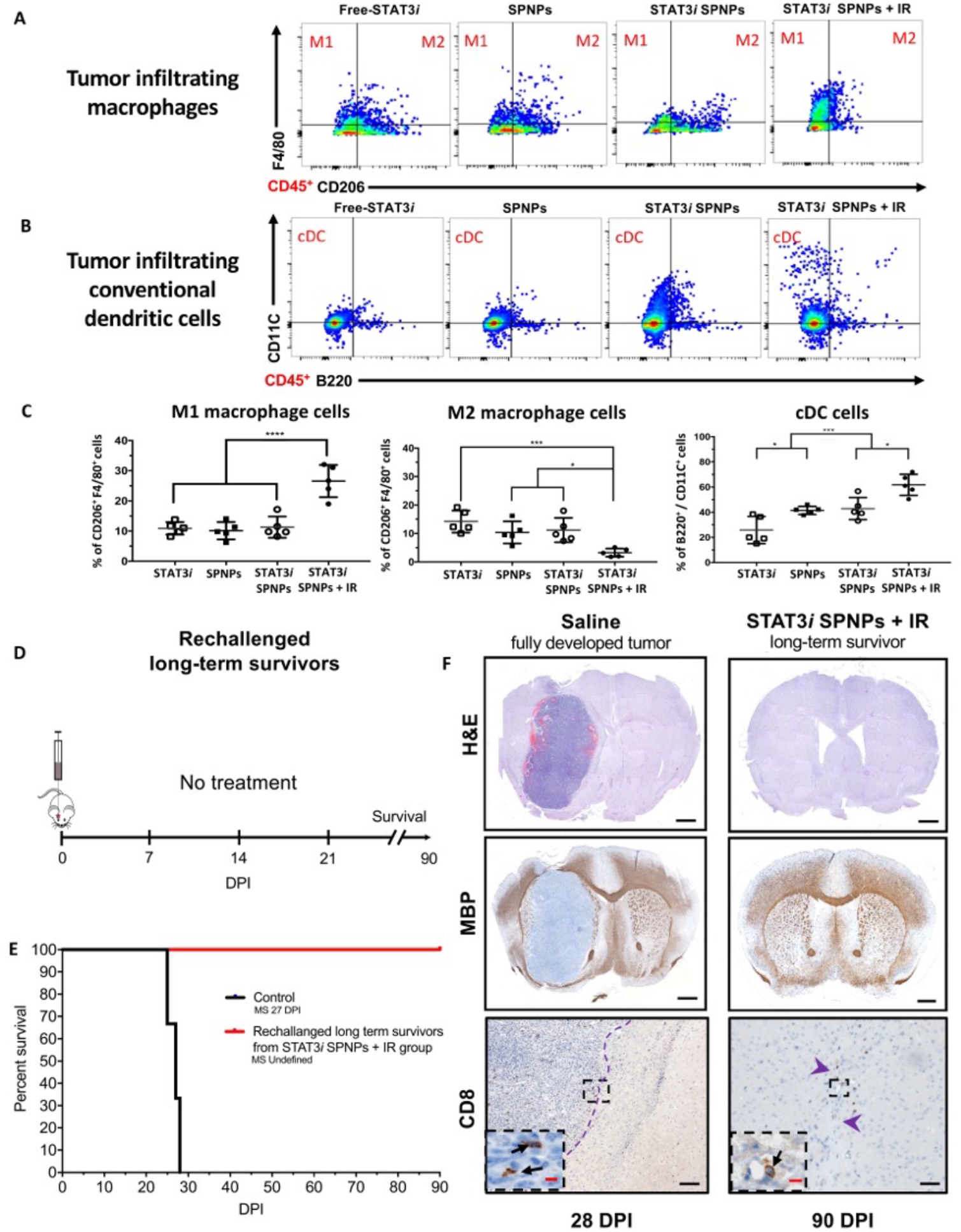
SPNPs protect against GBM rechallenge. **(A-B)** Flow analysis of tumor infiltrating **(A)** macrophage and **(B)** conventional dendritic cell (cDCs) populations in the tumor microenvironment (TME) following NP + IR treatment regimen. **(C)** Quantitative analysis of the immune cellular infiltrates showed a shift in the relative macrophage (M1*vs.* M2) present in the TME. In the free siRNA, empty SPNPs, and STAT3*i* SPNPs groups no significant change in the macrophage population were observed. Conversely, STAT3*i* SPNPs + IR treatment induces both a surge of approximately 2.5-fold increase in M1 population and a sharp 3- to 4-fold decrease in M2 macrophages. Among cDCs, progressively larger numbers of the cell population were observed moving from free siRNA to SPNPs groups. The combined treatment of STAT3*i* SPNPs with IR displayed the highest number of cDCs in the brain TME. **(D)** Timeline for rechallenging the long-term survivor for STAT3*i* SPNPs + IR survival study rechallenged, where following tumor implantation, no further treatment was provided. **(E)** Kaplan-Meier survival curve shows all rechallenged survivors reach a second long-term survival timepoint of 90 DPI in the absence of any therapeutic interventions. **(F)** IHC staining (H&E, MBP, and CD8) comparing the brains of untreated and rechallenged long-term survivors. No overt signs of remaining tumor, necrosis, inflammation, or disruption of normal brain architecture was observed in rechallenged long-term survivors from STAT3*i* SPNPs treatment group. Representative flow plots for each group are displayed. ****p < 0.0001; unpaired t-test. Bars represent mean ± SEM (n = 5 biological replicates).

## Tumor Re-challenge Study

Current standard of care approaches, including surgical resection combined with focused radiation and the chemotherapeutic temozolomide, have been used to treat primary GBM tumors. However, owning to the aggressive and infiltrative nature of GBM, these patients, as a rule, experience recurrence contributing to the high mortality and dismal survival rates. Based on the encouraging immune response observed in our survival study, we chose to re-challenge survivors from the STAT3*i* SPNP + IR treatment group. Tumors were implanted in the contralateral hemisphere of mice that were previously cured by the STAT3*i* SPNP + IR therapy. These mice did not receive any additional intervening therapy (Fig. 4d). As a control, naïve mice were also implanted with tumors at the same timepoint and likewise received no treatment. As expected, the control group saw rapid tumor growth, increased signs of disease and had a median survival of 27 DPI. Despite not receiving any additional treatment, all re-challenged mice survived to a second long-term survival point of 90 DPI (relative to the second tumor implantation, 180 days post initial tumor implantation) (Fig. 4e). IHC analysis of the brains yielded comparable results (Fig. 3f). H&E staining clearly showed the formation of a fully developed tumor mass in the control group, while members of the re-challenged cohort displayed no regions of necrosis, palisades or hemorrhages in either hemisphere (Fig. 4f, top). MBP staining confirmed that there was no overt disruption of the surrounding brain architecture (Fig. 4f, middle). Lastly, the presence of CD8 T cells was observed to be five-fold lower (Supplemental Fig. 15) in the STAT3*i* SPNP re-challenge group compared to the control (Fig. 4f, bottom). Importantly, we found no adverse effects in the brains of re-challenged survivors. Our findings suggest the potential involvement of an adaptive immune response that appears to guard against secondary tumors; an essential condition of any successful GBM therapy which will require long-term eradication of migrating and resistant CSCs, typically missed by traditional therapies.

## Conclusions

The main clinical challenge of GBM therapy is the lack of effective delivery systems that provide long-lasting health benefits for patients. A wide range of experimental approaches have been proposed in recent years based on either systemic or intracranial delivery strategies. To date, experimental studies that report even modest levels of efficacy are scarce and almost exclusively require invasive convection-enhanced intracranial delivery.^18,25^ As for systemic delivery, the situation is even more troublesome with only a couple of studies reporting long-term survivors at all.^12,29^ SPNPs combine the biological benefits of proteins with the precise engineering control of synthetic nanoparticles to yield (i) high efficacy (87.5% long-term survivors in a very aggressive intracranial tumor model), (ii) effective tumor delivery using systemically administered nanoparticles, and (iii) novel possibilities towards long-term eradication of resistant cancer cells using immunomodulatory protein nanoparticles. While protein-based nanoparticles have been a fairly uncharted area of research, this study demonstrates that synthetic nanoparticles that use proteins as structural building blocks may pave a novel route towards clinical cancer therapy. We studied the nanoparticle-mediated delivery of a siRNA against STAT3 (STAT3*i*), but the SPNP platform could be adopted, with further work, for delivery of small-molecule drugs, other siRNA therapies, or even drug combinations to a wide variety of solid tumors.

