## Supplemental Fig. 1 for "Systemic Brain Tumor Delivery of Synthetic Protein Nanoparticles for Glioblastoma Therapy"

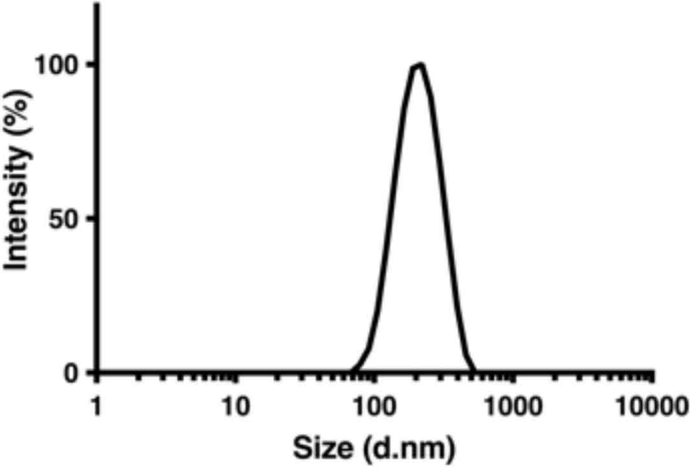


**Supplementary Fig. 1: SPNPs are stable at physiological conditions.** Characterization of SPNPs size was measured by dynamic light scattering (DLS) following the collection process and storage in PBS (pH 7.4) at 4°C. Particles were observed to swell with an increase in average size compared to their dry, crosslinked state. Average diameter = 220 ± 26.1 nm. No change was observed in average particle diameter over a period of 1 month at above conditions.
