## Supplemental Fig. 2 for "Systemic Brain Tumor Delivery of Synthetic Protein Nanoparticles for Glioblastoma Therapy"

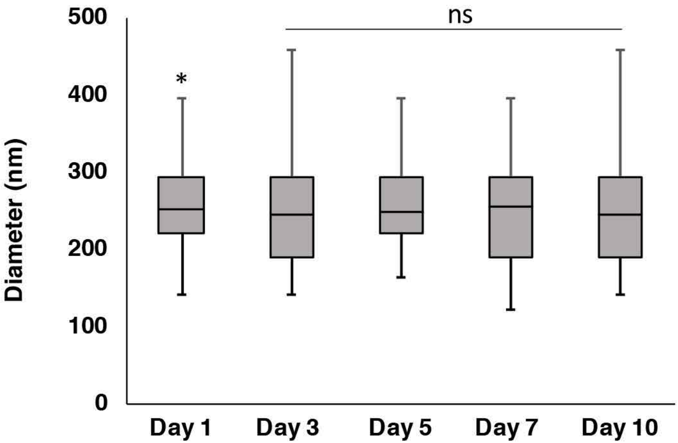


**Supplementary Fig. 2: SPNPs remain stable in solution under relevant physiological conditions.** After a single day in PBS at 37°C, SPNPs show no significant change in particle size as measured by dynamic light scattering (DLS). Particles appear to both remain intact and do not aggregate under these conditions. *$p < 0.05$
