## Supplemental Fig. 3 for "Systemic Brain Tumor Delivery of Synthetic Protein Nanoparticles for Glioblastoma Therapy"

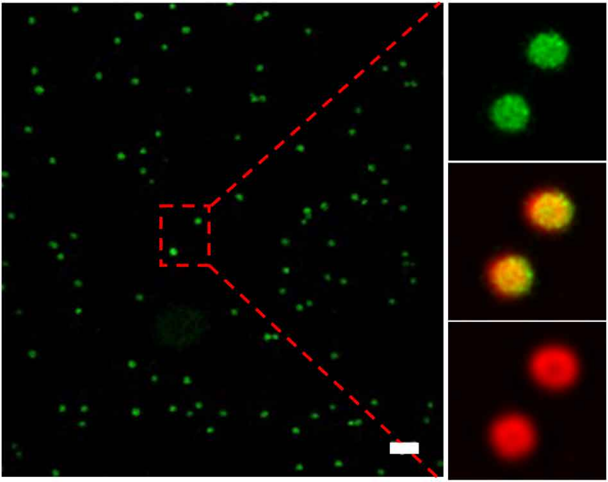

**Supplementary Fig. 3: SPNPs loaded with Cyanine3-siRNA exhibit controlled siRNA release.** Alexa Fluor 488 (green) labeled SPNPs loaded with a fluorescently (Cy3, red) labeled, scrambled siRNA. Imaging was performed using super-resolution, Stimulation Emission Depletion (STED) microscopy, which confirmed the localization of siRNA within the particles. Scale bar = 1 $\mu$m.
