## Supplemental Fig. 4 for "Systemic Brain Tumor Delivery of Synthetic Protein Nanoparticles for Glioblastoma Therapy"

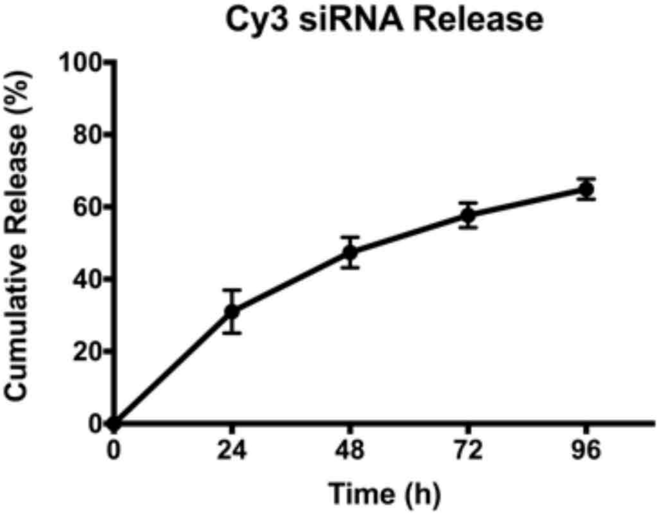


**Supplementary Fig. 4: Controlled siRNA release from SPNPs.** Loading and subsequent release of Cy3-labeled scrambled siRNA demonstrates a controlled and extended release of incorporated NP cargo. While 60% of the initially encapsulated siRNA is released in the first 96 hours at pH 7.4 and 37°C, continued release is observed for up to 21 days. Complete release confirms a loading efficiency of 96%.
