## Supplemental Fig. 5 for "Systemic Brain Tumor Delivery of Synthetic Protein Nanoparticles for Glioblastoma Therapy"

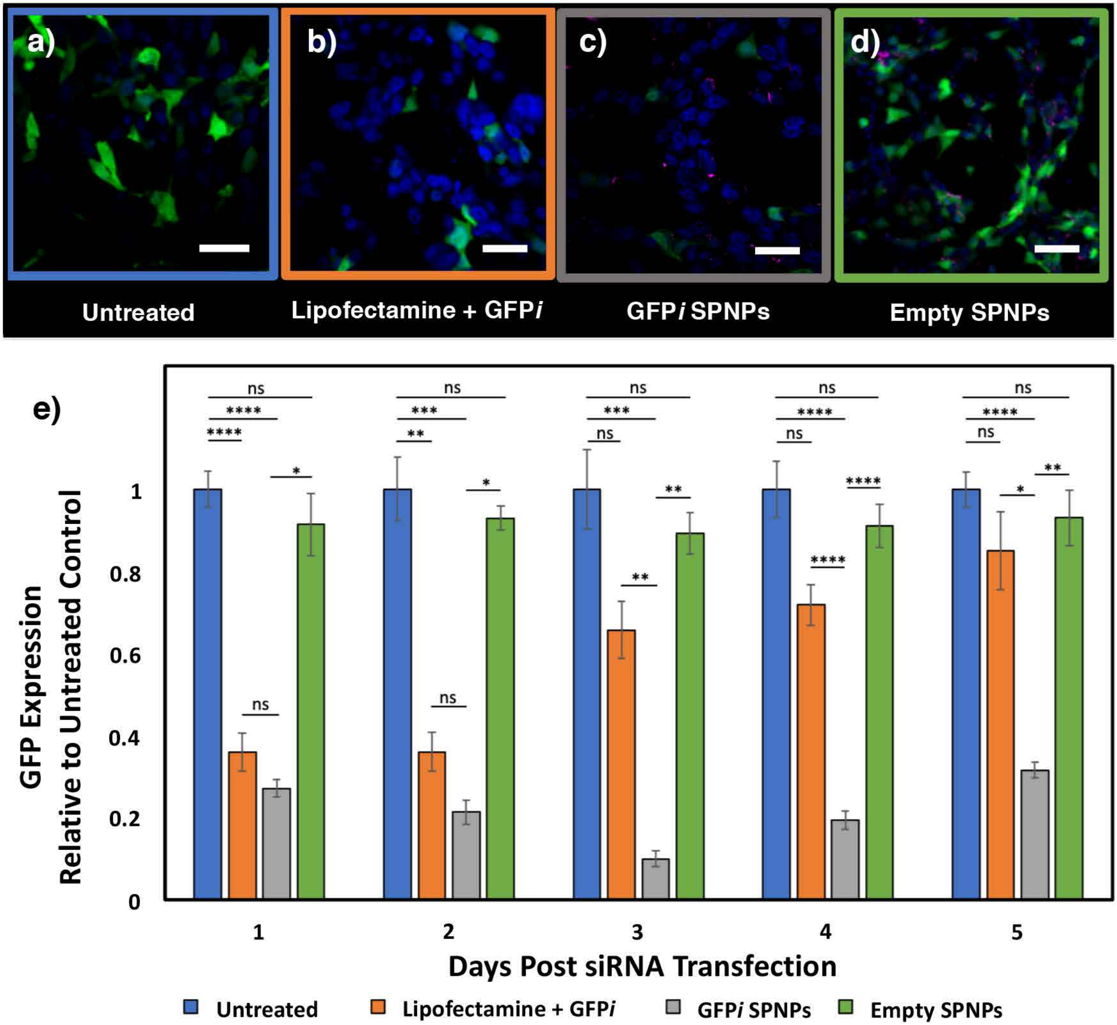


**Supplementary Fig. 5: *in vitro* GFP siRNA-loaded SPNPs reduce protein expression over a prolonged period. a-d,** Representative Confocal Scanning Laser Microscopy (CSLM) images of GL26-Cit cells incubated with NPs at 48-hour time point. **a,** Control group receiving no treatment. **b,** Positive control group transfected with GFP siRNA (GFP*i*) using Lipofectamine 2000. **c,** Cells treated with GFP*i* loaded nanoparticles at a concentration of 25 $\mu$g NPs/mL. **d,** Cells treated with empty albumin nanoparticles. **e**, GFP expression plotted relative to untreated control group over a period of 5 days. A significant and prolonged suppression of target protein is observed in cells that received the siRNA-loaded nanoparticles. A similar knockdown was observed in cells transfected with free siRNA using Lipofectamine at early time points, but a rapid recovery was observed after 48h. Bars represent the mean relative to untreated control ± standard error (n=3). ****$p < 0.0001$, ***$p < 0.0005$, **$p < 0.005$, *$p < 0.05$, ; unpaired t-test. Scale bars = 50 $\mu$m.
