## Supplemental Fig. 6 for "Systemic Brain Tumor Delivery of Synthetic Protein Nanoparticles for Glioblastoma Therapy"

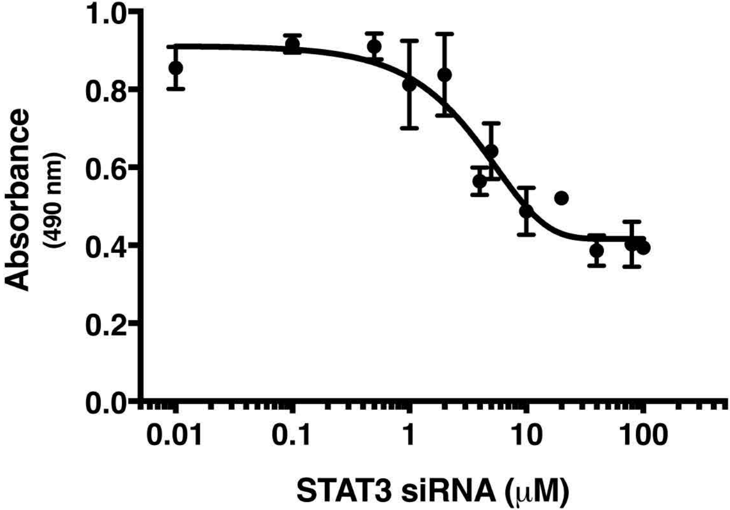


**Supplementary Fig. 6: STAT3 siRNA activity is non-toxic towards glioma cells.** Testing toxicity associated with free STAT3i, no toxicity was observed at relevant therapeutic concentrations. Concentrations delivered via SPNPs showed a significant silencing ability ($6.5 \times 10^{-4}$ $\mu$M), which were 6,000x less than the IC50 (3.85 $\mu$M) for soluble siRNA delivered via traditional transfection.
