## Supplemental Fig. 7 for "Systemic Brain Tumor Delivery of Synthetic Protein Nanoparticles for Glioblastoma Therapy"

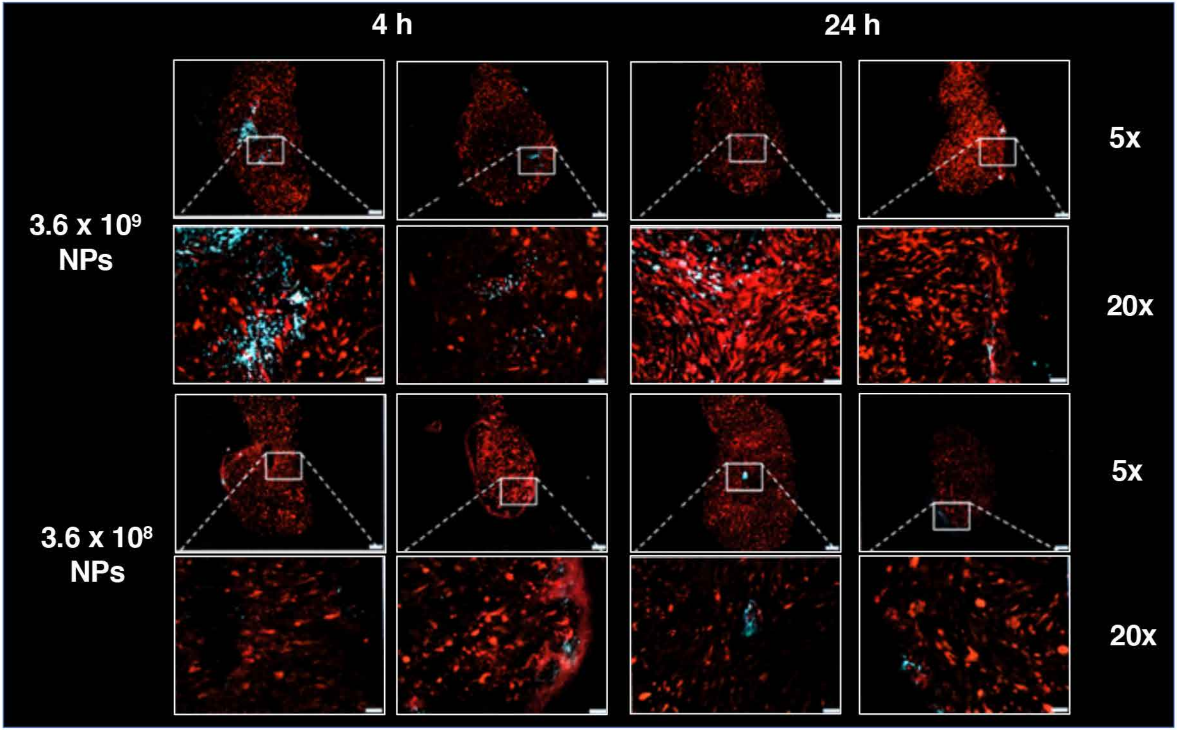


**Supplementary Fig. 7: SPNPs distribution in the tumor mass following intracranial injection.** Following the implantation of m-Tomato expressing GL26 tumors, C57BL/6 mice received 3 $\mu$L intracranial injections of SPNPs at either 3.6 x $10^8$ or 3.6 x $10^9$ per mL. Images suggest that the particles actively and rapidly distribute throughout the tumor mass. Scale bars = 50 $\mu$m.
