## Supplemental Fig. 8 for "Systemic Brain Tumor Delivery of Synthetic Protein Nanoparticles for Glioblastoma Therapy"

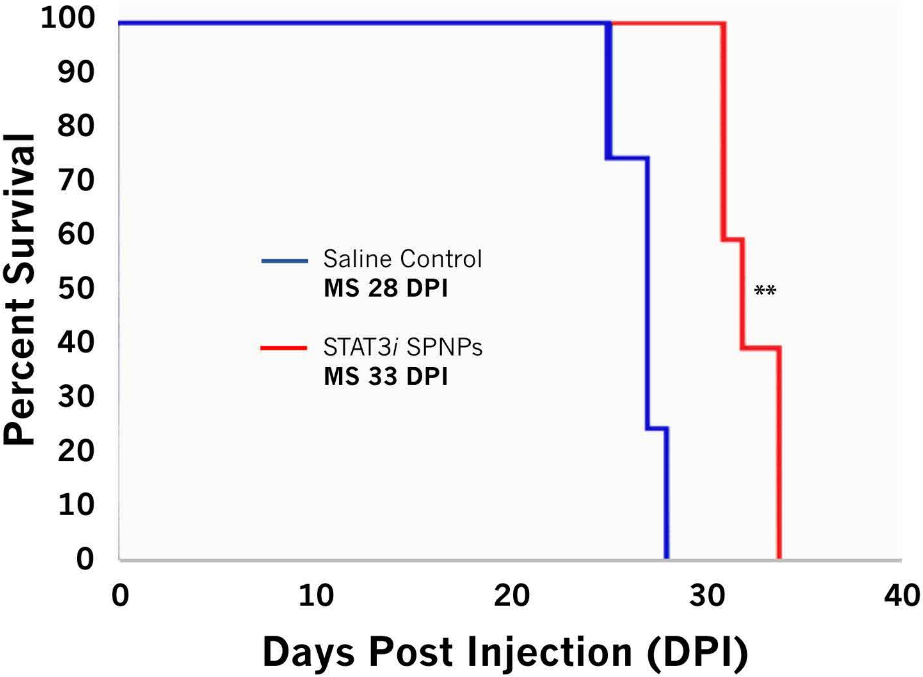


**Supplementary Fig. 8: Kaplan-Meier survival curve for single dose of systemically administrated STAT3i SPNPs.** C57BL6 mice were implanted with GL26 cells. A single dose of 2.0 x $10^{11}$ STAT3*i* SPNPs were delivered via tail vein injection five days post tumor implantation. Mice treated with siRNA-loaded particles had a median survival of 33 days, 5 days longer than mice in the saline treated control group (Data were analyzed using the log-rank (Mantel-Cox) test. $^{**}p < 0.001$ MS=median survival).
