## Supplemental Fig. 9 for "Systemic Brain Tumor Delivery of Synthetic Protein Nanoparticles for Glioblastoma Therapy"

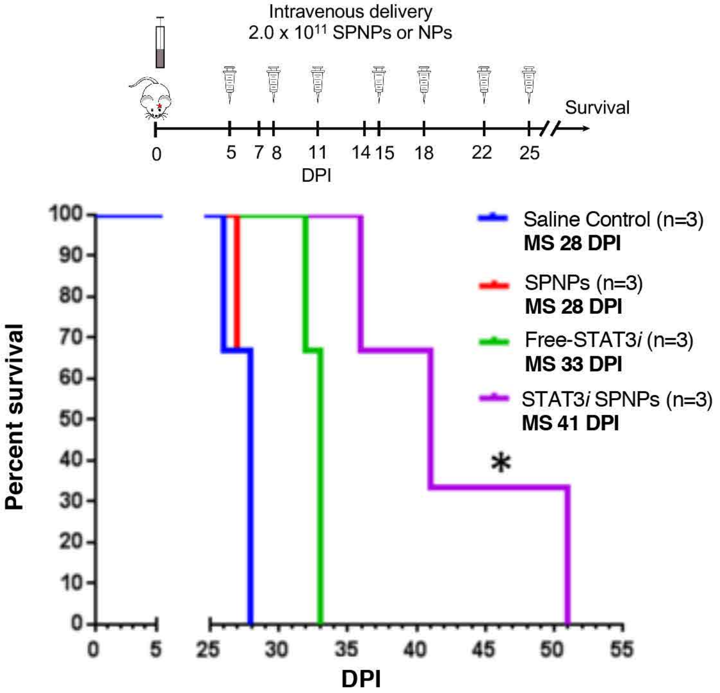


**Supplementary Fig. 9: Three week STAT3*i* SPNP treatment regimen extends median survival.** C57BL/6 mice were implanted with GL26 cells, at 5 DPI mice were treated with 2 x $10^{11}$ STAT3*i* SPNPs. STAT3*i* SPNPs delivered via tail vein (IV) injection. STAT3*i* SPNPs elicited a 46% increase in median survival compared to saline treated control mice (MS =41 vs. 28). Soluble IV administered STAT3*i* showed a moderate therapeutic effect (MS = 33 vs. 28) while mice treated with vehicle SPNPs saw no effect. (Data were analyzed using the log-rank (Mantel-Cox) test. *$p < 0.05$ MS=median survival).
