## Supplemental Fig. 10 for "Systemic Brain Tumor Delivery of Synthetic Protein Nanoparticles for Glioblastoma Therapy"

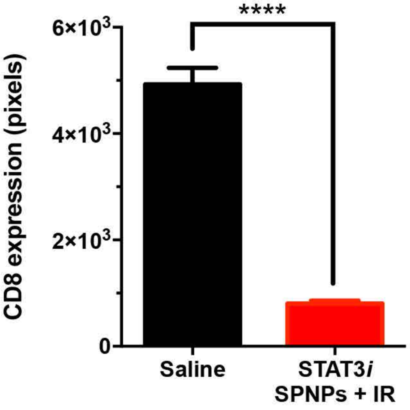


**Supplementary Fig. 10: Quantification of CD8 Expression in TME (90 DPI long-term survivor vs. 28 DPI control mouse at moribund stage).** Immunofluorescence staining in n = 3 different tumors in either saline or STAT3i TPNP + IR treatment groups was quantified using otsu threshold by ImageJ. Bar graphs represent total number of positive cells for CDS and in saline (28 dpi) vs. STAT3*i* SPNPs + IR (90 DPI) long-term survivor. ****p < 0.0001; unpaired t-test. Bars represent mean ± SEM (n = 3 biological replicates).
