## Supplemental Fig. 11 for "Systemic Brain Tumor Delivery of Synthetic Protein Nanoparticles for Glioblastoma Therapy"

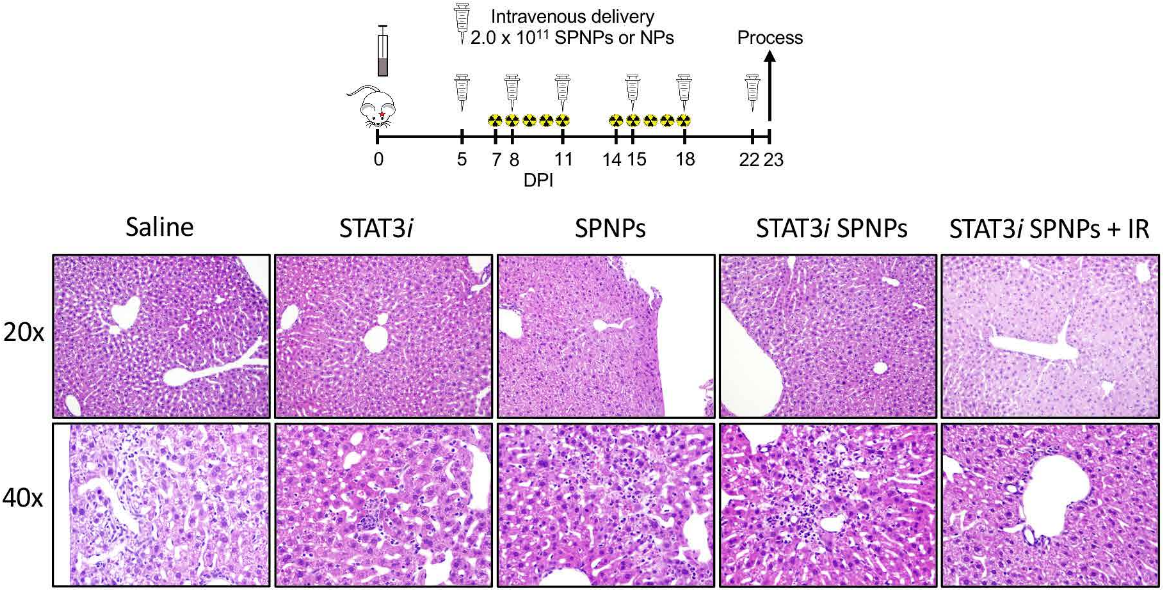


**Supplementary Fig. 11: STAT3*i* SPNPs and IR treatment do not impair liver function.** Histology performed on resected livers following a complete treatment of GMB tumor bearing mice find isolated regions of mild coagulative necrosis deemed to be well-contained and therefore would not induce a biological effect on liver function. In all groups , with the exception of the saline treated control, signs of hepatocellular necrosis was observed. This is attributed to water or glycogen accumulation in hepatocytes associated with a change in energy balance rather than a degenerative change.
