## Supplemental Fig. 12 for "Systemic Brain Tumor Delivery of Synthetic Protein Nanoparticles for Glioblastoma Therapy"

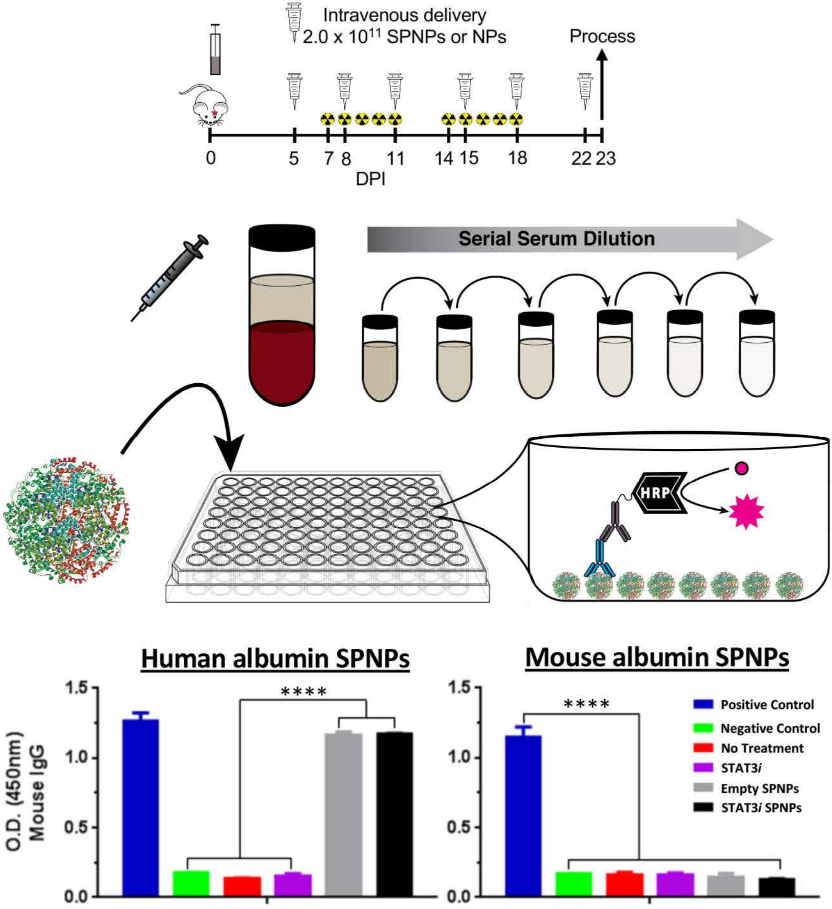


**Supplementary Fig. 12: Production of circulating antibodies against STAT3*i* SPNPs.** Following complete STAT3*i* SPNPs + IR treatment, circulating antibodies against human serum albumin nanoparticles were observed in serum of GL26 tumor bearing mice. Exchanging mouse serum albumin in the formulation, while maintaining all other components, eliminates this observed response. ****$p < 0.0001$; unpaired t-test. Bars represent mean ± SEM (n = 3 biological replicates).
