## Supplemental Fig. 13 for "Systemic Brain Tumor Delivery of Synthetic Protein Nanoparticles for Glioblastoma Therapy"

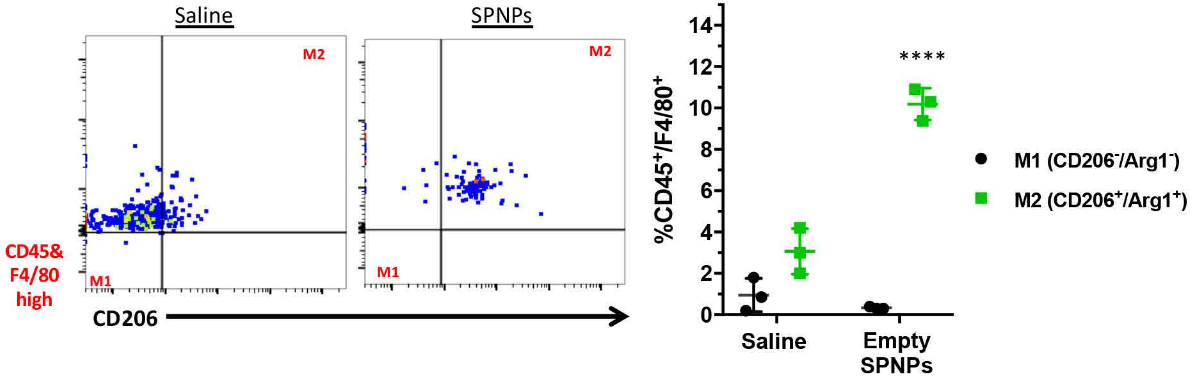


**Supplementary Fig. 13: SPNPs treatment induces a shift in the TME macrophage balance.** Treatment of GL26 tumor-bearing mice with Empty TPNPs produced a shift in macrophage populations within the tumor microenvironment. An increase in the M2 macrophages relative to saline treated control animals was observed. (n = 3 biological replicates, student t-test, $^{****}p<0.0001$)
