## Supplemental Fig. 14 for "Systemic Brain Tumor Delivery of Synthetic Protein Nanoparticles for Glioblastoma Therapy"

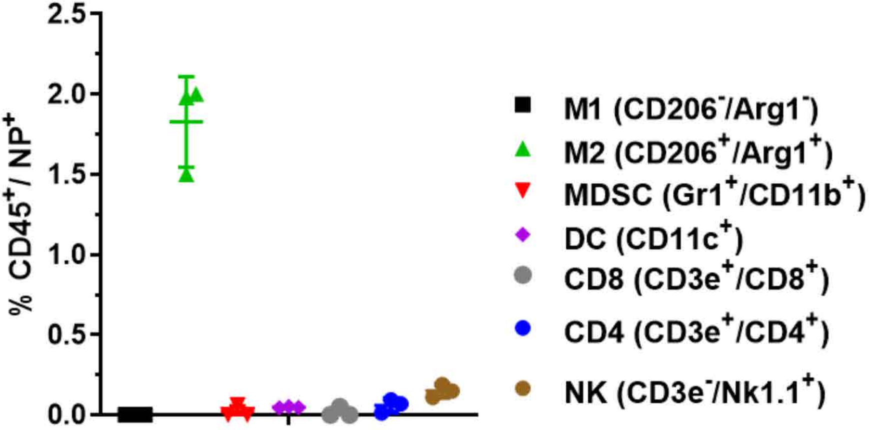


**Supplementary Fig. 14: Among TME immune cells, only M2 macrophages showed significant uptake of SPNPs**. Flow cytometry analysis of immune cells collected from the tumor microenvironment of SPNPs treated mice show significant nanoparticle uptake by M2 macrophages and minimal uptake by all other immune cell types (n = 3 biological replicates).
