## Supplemental Fig. 15 for "Systemic Brain Tumor Delivery of Synthetic Protein Nanoparticles for Glioblastoma Therapy"

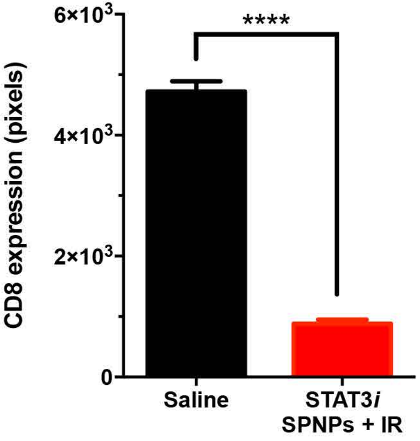


**Supplementary Fig. 15: Quantification of CD8 expression in TME (90 DPI rechallenged long-term survivor (180 days post initial tumor implantation vs. 28 DPI control mouse at moribund stage).** Immunofluorescence staining in n = 3 different tumors in each treatment group was quantified using otsu threshold by ImageJ. Bar graphs represent total number of positive cells for CD8 in saline (28 DPI) vs. STAT3*i* SPNPs + IR (60 DPI post rechallenged; 160 DPI post initial tumor implantation) rechallenged long term survivor ****$p < 0.0001$; unpaired t-test. Bars represent mean ± SEM (n = 3 biological replicates).
